# In Vivo Tensor-Valued Diffusion MRI Reveals Isotropic-Anisotropic Kurtosis Mismatch in a Rat MCAO Stroke Model

**DOI:** 10.1101/2025.06.23.661205

**Authors:** Mingyao Liang, Jiangyu Yuan, Chaogang Tang, Pingfu Wang, Tingting Gu, Zhao-Yu Deng, Weiming Sun, Xinming Yang, Yaohui Tang, Ye Li, Yi He

**Affiliations:** The Fifth Affiliated Hospital, Sun Yat-sen University, Zhuhai 519000, China; State Key Laboratory of Biomedical Imaging Science and System, Shenzhen Institute of Advanced Technology, Chinese Academy of Sciences, Shenzhen 518055, China; Paul C Lauterbur Research Center for Biomedical Imaging, Shenzhen Institute of Advanced Technology, Chinese Academy of Sciences, Shenzhen 518055, China; Department of Biomedical Engineering, Shanghai Jiaotong University, Shanghai 200030, China; Shenzhen University of Advanced Technology, Shenzhen 518107, China; National Innovation Center for Advanced Medical Devices, Shenzhen 518100, China

**Keywords:** Anisotropic-Isotropic Mismatch, Middle Cerebral Artery Occlusion, Ischemic Stroke, Tensor-Valued Diffusion MRI, Diffusion Kurtosis Imaging (DKI)

## Abstract

**Background:** Tensor-valued diffusion MRI (dMRI) enables the separation of total kurtosis into isotropic and anisotropic components, offering improved specificity over conventional Diffusion Kurtosis Imaging (DKI). In this study, we introduce a novel framework for detecting anisotropic-isotropic kurtosis mismatch and evaluate its utility in characterizing stroke lesions.

**Methods:** We performed tensor-valued dMRI in a rat model of middle cerebral artery occlusion (MCAO). Metrics including anisotropic kurtosis (MK_A_) and isotropic kurtosis (MK_I_) were quantified to assess microstructural tissue heterogeneity. Histological validation was conducted using co-registered tissue sections.

**Results:** We observed significant mismatches between mean diffusivity (MD), mean kurtosis (MK), as well as specifically in MK_A_ and MK_I_ in ischemic regions. Notably, MK_A_ and MK_I_ exhibited statistically significant alterations compared to contralateral regions. Moreover, these metrics strongly correlated with pathological findings. The mismatch highlighted microstructurally distinct subregions within the lesion.

**Conclusions:** This is the first study to propose an anisotropic-isotropic kurtosis mismatch framework in vivo. Tensor-valued dMRI may offer a more specific imaging biomarker for stroke, with potential for improving lesion subtyping and guiding therapeutic decisions.

## 1. Introduction

Ischemic stroke is a leading cause of disability and death worldwide, underscoring the need for precise neuroimaging to support early diagnosis, guide therapy and improve patient outcomes^1, 2^. Diffusion-weighted MRI (DWI) has revolutionized acute stroke assessment by enabling early detection of ischemic lesions within minutes of symptom onset^3, 4^. Conventional diffusion tensor imaging (DTI), which derives metrics such as mean diffusivity (MD) and fractional anisotropy (FA), provides insights into water mobility and directional tissue organization. However, DTI relies on the assumption of Gaussian diffusion, limiting its sensitivity to the complex microstructural changes after stroke^5^. While diffusion restriction (manifested as decreased MD) indicates cytotoxic edema in acute ischemia, DTI cannot fully resolve tissue heterogeneity or distinguish reversible injury^6^.

To overcome these limitations, advanced diffusion MRI techniques have been developed to characterize ischemic tissue beyond the capabilities of DTI^5, 7^. Diffusion kurtosis imaging (DKI), an extension of DTI, quantifies non-Gaussian diffusion of water and is more sensitive to microstructural heterogeneity^3, 8^. DKI has demonstrated improved sensitivity in detecting acute ischemic lesions compared to DTI^7^. Early studies reported elevated kurtosis values in regions of acute infarction, offering additional contrast between ischemic and normal brain tissue^9^. Specifically, mean kurtosis (MK) is typically increased in infarcted areas relative to the contralateral hemisphere, while mean diffusivity (MD) is decreased. Notably, infarct volumes defined by MK often appear smaller than those delineated by DWI or MD, suggesting that kurtosis imaging may more precisely identify the damaged infarct core within the broader diffusion-restricted region^10^.

Despite its promise, the biological interpretation of kurtosis contrast remains challenging. Kurtosis can arise from several microstructural features, including tissue anisotropy (variations in diffusion due to oriented structures), heterogeneity of diffusivities across compartments, and microscopic diffusion restrictions or disorder at the cellular level^7^. Standard DKI collapses all these sources into a single MK metric, obscuring their individual contributions and limiting biological specificity. To address this limitation, tensor-valued diffusion MRI has emerged as a powerful technique that leverages advanced b-tensor encoding schemes^11–13^. Unlike conventional linear encoding (single-direction), tensor-valued encoding employs multiple b-tensor shapes (such as linear, planar, and spherical) to probe diffusion properties in a multidimensional manner^14, 15^. This multidimensional approach enables the decomposition of total kurtosis into anisotropic kurtosis (MK_A_), which reflects orientation-dependent microstructural features such as axonal coherence or dispersion, and isotropic kurtosis (MK_I_), which captures orientation-independent restrictions caused by phenomena such as cell swelling or high cell density^16, 17^. Histologically, MK_A_ corresponds to histological anisotropy (H_A_) on the microscopic scale, influenced by the eigenvalues of the local structure tensor. In contrast, MK_I_ captures histological isotropy (H_I_) on the microscopic scale, related to variable cell densities^14, 15^. This decomposition offers a more nuanced view of tissue microstructure, potentially enhancing the specificity of diffusion MRI in pathological conditions.

In the context of acute ischemic stroke, the brain can be divided into a damaged infarct core and a surrounding ischemic penumbra, which is functionally impaired but potentially salvageable^18–20^. Two major imaging mismatch paradigms are widely recognized: (1) Perfusion-Diffusion Mismatch (PWI-DWI mismatch)^21–23^: a macroscopic concept identifying areas with reduced blood flow but preserved tissue integrity. (2) Diffusivity-Kurtosis Mismatch^10^ (MD-MK mismatch): A microscopic concept based on diffusion kurtosis imaging to assess microstructural damage. In particular, a mismatch between mean diffusivity (MD) and mean kurtosis (MK) has been proposed to delineate damaged core tissue from potentially salvageable penumbra^10^. However, a significant limitation of DKI is its lack of specificity, its aggregate MK measure cannot distinguish between distinct microstructural alterations^24^.

To our knowledge, no in vivo study has yet applied tensor-valued diffusion MRI in a rodent model of cerebral ischemia to explicitly quantify the mismatch between anisotropic and isotropic kurtosis. Identifying this kurtosis mismatch may enhance the delineation of injured tissue and provide a novel imaging biomarker for stroke. In this study, we addressed this gap by applying tensor-valued dMRI in a rat model of MCAO, closely resembles the pathophysiological features of heterogeneous tissue injury in humans^3, 25^. We hypothesized that in vivo quantification of anisotropic kurtosis (MK_A_) and isotropic kurtosis (MK_I_) would sensitively detect microstructural alterations within ischemic brain tissue and reveal a measurable mismatch between MK_A_ and MK_I_ within the lesion. We further evaluated the diagnostic utility of this mismatch in comparison with standard DKI metrics MK and DTI metric MD, and correlated imaging findings with histopathological markers of tissue injury. Our ultimate goal is to establish whether tensor-valued dMRI can provide a more specific imaging biomarker, particularly the anisotropic-isotropic kurtosis mismatch for stroke characterization, with the potential to guide treatment decisions and improve prognostication.

## 2. Methods

### 2.1. Animal Model

All animal procedures were conducted in accordance with a protocol approved by the Animal Ethics Committee of The Fifth Affiliated Hospital, Sun Yat-sen University (Protocol No. 00328) and adhered to institutional and national guidelines for laboratory animal care and use. Male Sprague-Dawley rats (n = 6; 8-12 weeks old) were housed under a 12-hour light/dark cycle with ad libitum access to food and water. A middle cerebral artery occlusion (MCAO) rodent model was used in the study^26, 27^. Cerebral ischemia was induced using the intraluminal filament model of middle cerebral artery occlusion (MCAO). A silicone-coated monofilament was inserted into the internal carotid artery via the external carotid artery and advanced to occlude the origin of the middle cerebral artery. After 2 hours of occlusion, the filament was withdrawn to allow reperfusion, thus establishing a reproducible model of transient ischemia-reperfusion injury.

### 2.2. MRI Acquisition and Data Processing

All MR images were collected using a 9.4T small animal MRI system (BioSpec 94/30, Bruker, Germany). Tensor-valued diffusion MRI (dMRI) acquisitions included both linear tensor encoding (LTE) and spherical tensor encoding (STE) schemes, with the following parameters: b-values = 200, 700, 1400, and 2000 s/mm^2^; Repetition time (TR): 6500 ms; Echo time (TE): 45 ms; Voxel Size: 250 x 250 x 250 μm^3^; Field of View (FOV): 32 × 32 mm^2^; Slices: 32 slices; number of segments = 8; and slice thickness = 800 µm. For standard DTI/DKI acquisition, the following diffusion scheme was used: b-values=1000 s/mm^2^ with 30 directions, b-values = 2000 s/mm^2^ with 60 directions, b = 0 s/mm^2^ with 5 non-diffusion weighted images.

Raw MRI data were processed and analyzed using classical software or programs in the field of MRI: MATLAB, FSL (https://fsl.fmrib.ox.ac.uk/fsl/fslwiki), MRtrix3 (https://www.mrtrix.org/), and Advanced Normalization Tools (ANTs). Imaging data exported from the MRI scanner in DICOM format were first converted to the NIfTI (Neuroimaging Informatics Technology Initiative) format, a standard for neuroimaging analysis. The NIfTI files were then converted to MRtrix Image Format (MIF) using MRtrix3, which enabled automated denoising to improve the signal-to-noise ratio (SNR). Subsequent preprocessing steps, including distortion correction, skull stripping, and spatial smoothing, were performed using FSL. Multidimensional diffusion parameter estimation was conducted with the md-dmri toolbox in MATLAB (https://github.com/markus-nilsson/md-dmri). Tensor-valued dMRI metric maps were generated, including mean diffusivity (MD), mean kurtosis (MK), mean isotropic kurtosis (MK_I_), and mean anisotropic kurtosis (MK_A_). Lesion areas were segmented using a k-means clustering algorithm applied to each diffusion map and then overlaid on corresponding T2-weighted anatomical images.

### 2.3. Histological Preparation

Following MRI acquisition, all animals were euthanized by transcardial perfusion with saline followed by 4% paraformaldehyde. Brains were post-fixed, paraffin-embedded, and sectioned at 5 μm thickness. Sections were stained with hematoxylin and eosin (HE) for general cytoarchitecture and Luxol fast blue (LFB) for myelin integrity assessment.

#### 2.3.1. Hematoxylin and Eosin (H&E) Staining

Paraffin sections were deparaffinized, rehydrated, and stained with hematoxylin for 5 minutes, rinsed under running tap water for 1 minute, followed by eosin staining for 5 minutes. Sections were dehydrated through graded ethanol, cleared in xylene for 10 minutes, and mounted with resin.

#### 2.3.2. Luxol Fast Blue (LFB) Staining

Sections were deparaffinized and rehydrated, then incubated in 0.1% Luxol fast blue solution at 60°C for 4 hours. After staining, sections were rinsed and differentiated in lithium carbonate for 5 seconds and then in 70% ethanol for 10 seconds. After thorough rinsing, slides were dehydrated in absolute ethanol (3 × 5 minutes), cleared in xylene for 5 minutes, and sealed with mounting resin.

### 2.4 Histology-MRI Co-registration and Region-of-Interest (ROI) Definition

Histological and MRI images were spatially co-registered using MRtrix3 and FSL software. High-resolution histological images (1.0 μm/0.1214 pixel) were aligned to the MRI data using custom-written MATLAB scripts. Image resolutions were normalized to ensure consistent scale across modalities. Identical ROIs were manually delineated on both histological and MRI datasets to enable quantitative comparison of microstructural features across imaging modalities.

### 2.5 Histological Quantification of histological isotropy (H_I_) and histological anisotropy (H_A_)

To validate microstructural parameters, H_I_ and H_A_ were quantitatively assessed from HE and LFB-stained tissue sections using in-house MATLAB algorithms. Each HE- and LFB-stained image was divided into 125×125 μm² regions, further subdivided into twenty-five 25×25 μm² subregions. H_I_ was calculated as the normalized variance of cell densities derived from microscopy^14^. Cell nuclei were segmented through grayscale thresholding and morphological operations, and the number of labeled nuclei was used to determine local cell density (cells/μm²). H_A_ was defined as the normalized variance of these eigenvalues, reflecting the degree of directional coherence^14^. LFB-stained images were converted to grayscale and analyzed using structure tensor analysis to quantify directional microstructural organization. Local image gradients were computed using Sobel operators, and the resulting tensor elements were smoothed with Gaussian filtering. Within each region, the dominant eigenvalues of the smoothed structure tensors were extracted. Regions of interest (ROIs) were manually defined based on tissue morphology following previously established protocols.

### 2.6 Statistical Analyses

All statistical analyses and data visualizations were performed using GraphPad Prism 10.0 (https://www.graphpad-prism.cn/) and MATLAB. Linear regression analyses were used to quantify the relationship between dMRI-derived parameters and histologically defined microstructural metrics, such as MK_A_ versus histological anisotropy (H_A_), and MK_I_ versus histological isotropy (H_I_). To evaluate group-level differences, two-sample t-tests were used to compare dMRI parameters between infarcted and contralateral brain regions. Pearson’s correlation coefficient (r) was used to assess the strength of associations, with statistical significance set at α = 0.05. *p* values < 0.05 were considered statistically significant. Statistically significant correlations are marked with asterisks (*), while non-significant correlations are denoted as “NS.”

## 3. Results

### 3.1 Induction of Middle Cerebral Artery Occlusion and Ischemia-Reperfusion Injury

We established a rat model of ischemic stroke using the intraluminal filament method for middle cerebral artery occlusion (MCAO), followed by reperfusion. High-resolution anatomical images were acquired using a 9.4T Bruker MRI system (**Figure 1A**). After in vivo imaging, animals were euthanized, and brains were harvested via transcardial perfusion for ex vivo analysis. Conventional diffusion MRI revealed microstructural alterations consistent with acute ischemia, including reduced mean diffusivity (MD) and elevated mean kurtosis (MK) within the affected hemisphere (**Figure 1B**). The spatial reproducibility and anatomical consistency of infarcts across animals were validated by both T2-weighted imaging and 2,3,5-triphenyltetrazolium chloride (TTC) staining, which confirmed lesion location and volume (**Figure 1C**).

**Figure 1.**
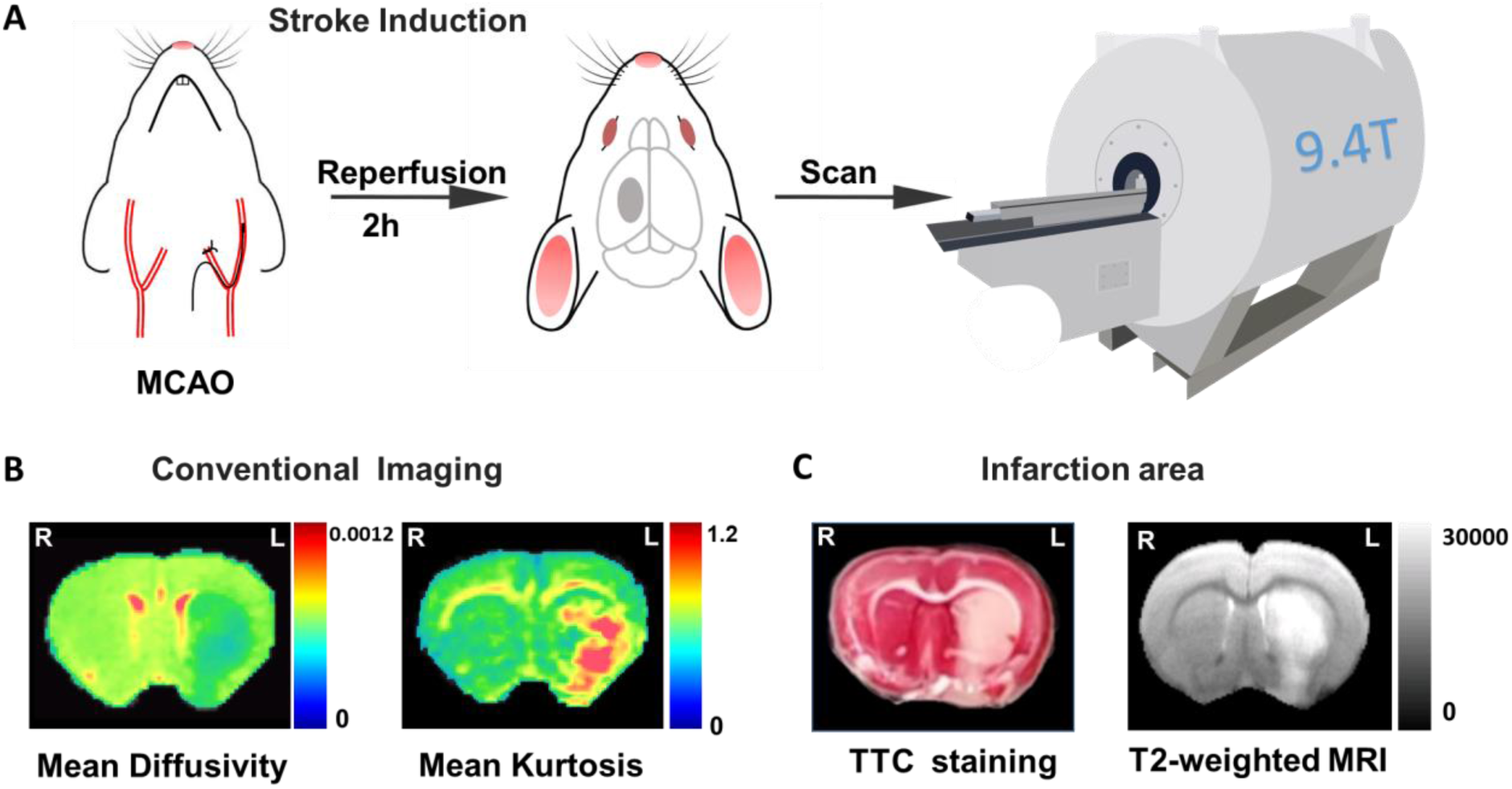
Experimental setup and validation of infarction in MCAO stroke model. **A,** Focal cerebral ischemia was induced using the middle cerebral artery occlusion (MCAO) model. Reperfusion was initiated 2 hours after occlusion, followed by imaging on a 9.4T MRI scanner. **B,** Conventional diffusion MRI maps show changes in tissue microstructure. Left: Mean diffusivity map. Right: Mean kurtosis map. **C,** Infarct localization and size were confirmed by TTC staining and T2-weighted imaging, demonstrating consistent lesion patterns across animals.

### 3.2 In Vivo Diffusion Encoding and Acquisition Using Tensor-Valued MRI in Stroke

To evaluate stroke-related tissue alterations, we performed both tensor-valued diffusion MRI and conventional diffusion weighted imaging in rats with MCAO-induced ischemia. The tensor-valued protocol included Linear Tensor Encoding (LTE) and Spherical Tensor Encoding (STE) acquisitions (**Figure 2A**), while conventional DWI employed standard single-direction diffusion encoding (**Figure 2B**). We analyzed powder-averaged signal decays from LTE and STE acquisitions in both the ipsilateral (lesion) and contralateral (normal) hemispheres. Orientation-invariant diffusion signals were obtained by averaging the diffusion-weighted volumes for each b-value shell, yielding metrics free from fiber orientation dispersion.

**Figure 2.**
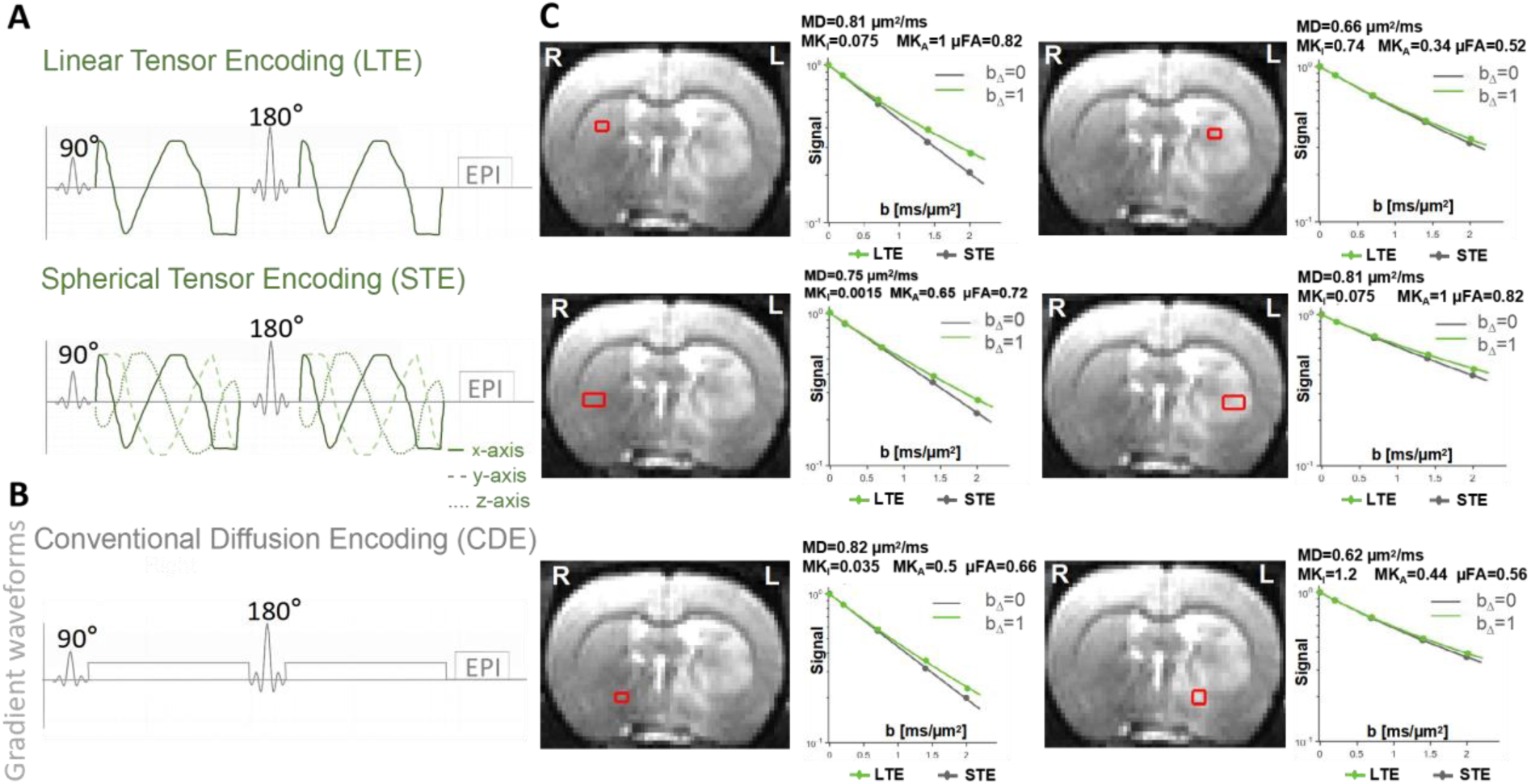
Tensor-valued diffusion MRI gradient waveforms and powder-averaged signal analysis. **A,** Gradient waveform schematics for linear tensor encoding (LTE) and spherical tensor encoding (STE), illustrating diffusion sensitization along single and multiple axes, respectively. **B,** Conventional pulsed gradient spin-echo (PGSE) waveform used in standard diffusion MRI. **C,** Representative b0 images showing regions of interest (ROIs) on the contralateral (normal) and ipsilateral (lesion) hemispheres. Corresponding powder-averaged diffusion signal decay curves from LTE (green) and STE (gray) are plotted, with derived metrics including mean diffusivity (MD), anisotropic and isotropic kurtosis (MK_A_, MK_I_), and microscopic fractional anisotropy (μFA).

Representative signal decays and derived microstructural metrics from six brain regions of interest (ROIs) in one MCAO rat are shown in **Figure 2C**. In the contralateral (non-ischemic) hemisphere, diffusion profiles showed higher MD values (e.g., 0.81-0.82 μm²/ms) and relatively lower MK_I_ values, with MK_A_ up to 0.5-1.0 and μFA around 0.66-0.82. In contrast, ischemic ROIs in the ipsilateral hemisphere demonstrated reduced MD values (e.g., as low as 0.62-0.66 μm²/ms), elevated MK_I_ (up to 1.2), and decreased MK_A_ and μFA values, in some cases as low as 0.34 and 0.52, respectively.

### 3.3 Regional Kurtosis Alterations Revealed by Tensor-Valued dMRI

In the MCAO rat model, comparison of the ipsilateral (lesion) and contralateral (non-lesioned) hemispheres revealed distinct alterations across tensor-valued diffusion MRI metrics. As shown in **Figure 3A**, ischemic regions exhibited markedly reduced mean diffusivity (MD) and anisotropic kurtosis (MK_A_), alongside elevated mean kurtosis (MK) and isotropic kurtosis (MK_I_). These patterns reflect both restricted water diffusion and increased microstructural complexity in the infarct core. Notably, MK_I_ maps highlighted regions with strong isotropic restriction, possibly corresponding to cytotoxic edema, inflammatory infiltration, and reactive gliosis, while MK_A_ was notably diminished in the same areas, likely reflecting disruption of coherent fiber architecture. The spatial mismatch between MK_A_ and MK_I_ further underscores the utility of tensor-valued decomposition in identifying microstructural heterogeneity within infarcts.

**Figure 3.**
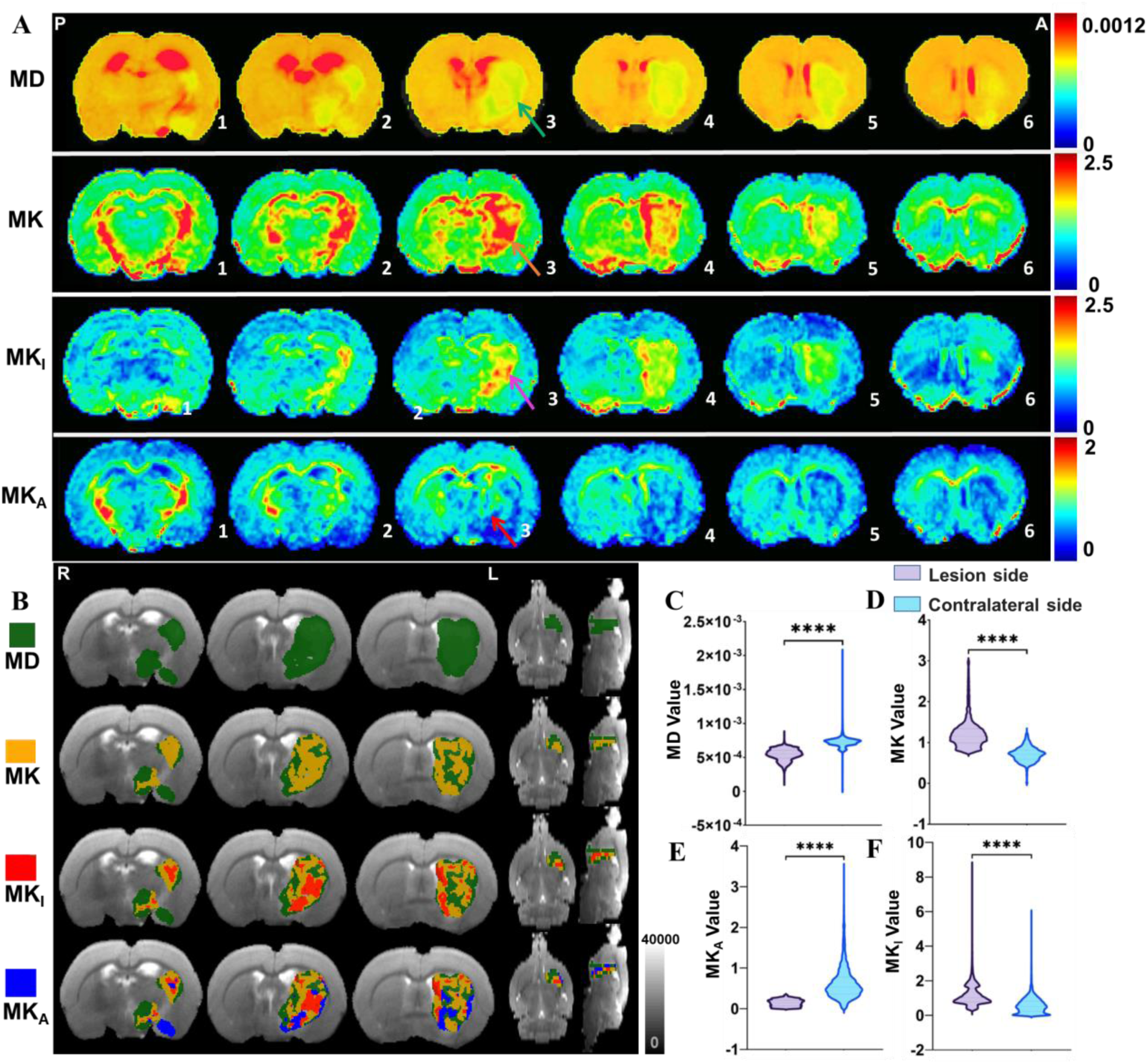
Mismatch between isotropic and anisotropic kurtosis in ischemic brain tissue. **A,** Representative coronal maps of mean diffusivity (MD), mean kurtosis (MK), isotropic kurtosis (MK_I_), and anisotropic kurtosis (MK_A_) from posterior (P) to anterior (A) brain slices. The lesion region demonstrates decreased MD (green arrow) and MK_A_ (red arrow), while MK (orange arrow) and MK_I_ (purple arrow) are notably reduced, indicating a mismatch between the kurtosis sources. **B,** Overlay of lesion regions defined by each diffusion metric (color-coded) on b0 images from a representative animal. ROIs were segmented based on metric-specific abnormal voxels: MD (green), MK (yellow), MK_I_ (red), and MK_A_ (blue). **C-F,** Voxel-wise comparisons between lesion and contralateral hemispheres for MD, MK, MK_A_, and MK_I_. All metrics show statistically significant differences (****, *p* < 0.0001, n=6 rats, two-sample t-test), supporting differential sensitivity to tissue microstructural changes.

To accurately delineate these variations, k-means clustering was applied to the multimetric data (**Figure S1**), enabling data-driven segmentation of infarcted regions based on voxelwise diffusion properties^3, 28^. We then overlaid lesion maps derived from each metric (**Figure 3B**), revealing that MK_I_-defined lesions were spatially more confined than MK-defined regions, while MK_A_ reductions extended variably across perilesional tissue. This provides further support for a kurtosis-based microstructural mismatch framework, where elevated MK may result from either increased isotropic restriction or retained directional coherence. Quantitative analysis of mean values across segmented lesion ROIs demonstrated statistically significant differences between hemispheres (**Figure 3C-F**). Compared to the contralateral side, lesion ROIs showed significantly lower MD and MK_A_ values (*p* < 0.0001), and significantly higher MK and MK_I_ values (*p* < 0.0001), confirming both diffusion restriction and kurtosis heterogeneity within ischemic tissue.

### 3.4 Histological Validation of Tensor-valued dMRI metrics in stroke

To validate diffusion MRI findings at the microstructural level, we performed region-of-interest (ROI) analysis by precisely co-registering in vivo dMRI maps with corresponding histological sections. Hematoxylin and eosin (HE) staining and Luxol fast blue (LFB) staining were used to identify cytological and myelin-specific changes, respectively, in infarcted versus contralateral brain tissue. For each dMRI-derived metric, symmetric ROIs were selected in both hemispheres to quantify anisotropic (MK_A_) and isotropic (MK_I_) kurtosis, alongside corresponding histological indices of structural integrity (H_A_ and H_I_). As shown in **Figure 4A** and **4C**, the infarcted regions demonstrated pronounced changes in both kurtosis metrics and their histological correlates.

**Figure 4.**
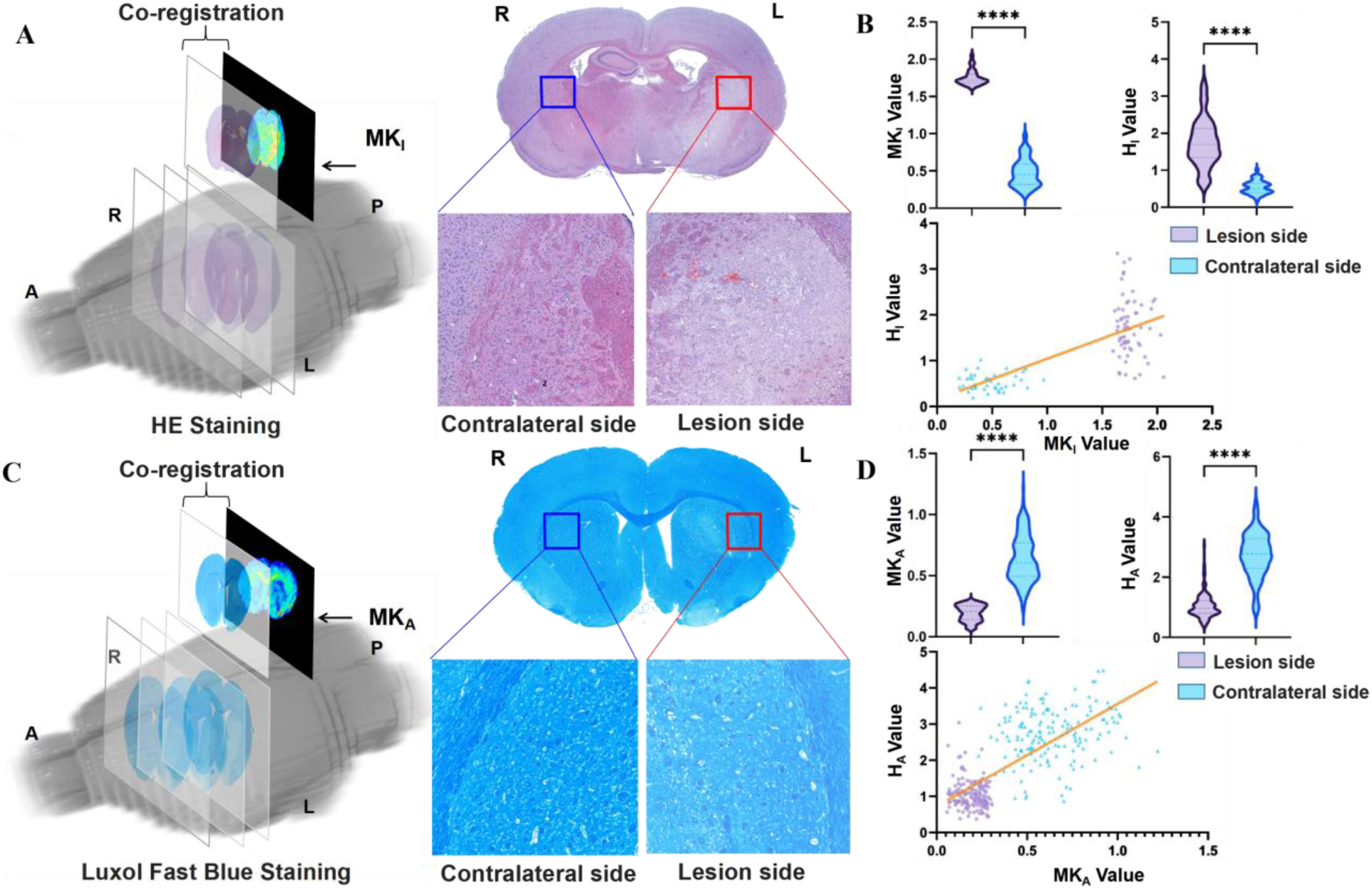
Histological validation of tensor-valued diffusion MRI metrics in ischemic brain tissue. **A,** Co-registration of MRI-derived isotropic kurtosis (MK_I_) maps with hematoxylin and eosin (HE) staining. Histological isotropy (H_I_) was assessed in the lesion and contralateral regions and compared with corresponding MK_I_ values. **B,** Violin plots show significantly elevated MK_I_ and histology-derived cell density index (H_I_) in the lesion side compared to the contralateral side (****, *p* < 0.0001). Linear regression reveals a strong positive correlation between MK_I_ and H_I_ (r = 0.7266, *p* < 10⁻⁵). **C**, Co-registration of anisotropic kurtosis (MK_A_) maps with Luxol Fast Blue (LFB) staining, used to quantify myelin density. **D,** MK_A_ and histology-derived anisotropy index (H_A_) were both significantly lower in the lesion side (****, *p* < 0.0001), and showed a strong positive correlation (r = 0.7246, *p* < 10⁻⁵), supporting MK_A_ as a marker of microstructural anisotropy loss.

Quantitative comparisons revealed significantly elevated MK_I_ and H_I_ values on the lesion side compared to the contralateral side (****, *p* < 0.0001, n = 6 rats; **Figure 4B**). Moreover, MK_I_ was strongly correlated with H_I_ (r = 0.7266, *p* < 10^−5^, n = 6 rats), suggesting that isotropic kurtosis reflects variable cell density induced by cytotoxic edema. Similarly, MK_A_ and H_A_ were significantly reduced in the infarcted hemisphere (****, *p* < 0.0001, n = 6 rats; **Figure 4D**), and a robust positive correlation was observed between MK_A_ and histological anisotropy H_A_ (r = 0.7246, *p* < 10^−5^, n = 6 rats). This supports the interpretation that anisotropic kurtosis is sensitive to loss of fiber organization and myelin architecture. Together, these data demonstrate that tensor-valued dMRI metrics, particularly MK_I_ and MK_A_, reliably correspond to histologically validated microstructural alterations in ischemic brain tissue.

### 3.5 Detection of Anisotropic–isotropic Mismatch by Tensor-Valued dMRI

Tensor-valued diffusion MRI enables the decomposition of mean kurtosis into isotropic (MK_I_) and anisotropic (MK_A_) components, providing enhanced specificity in identifying the microstructural origins of diffusion heterogeneity in ischemic tissue. This separation allows differential mapping of spherically restricted diffusion (e.g., cytotoxic edema) and directionally coherent structures (e.g., axons or dendrites). In **Figure 5A**, we reconstructed and visualized three-dimensional lesion masks based on each diffusion metric: MD, MK, MK_I_, and MK_A_. These reconstructions revealed distinct spatial distributions within the infarcted hemisphere. MK_I_-based lesions appeared more confined and medially located compared to the broader areas delineated by MD or MK. In contrast, MK_A_ highlighted regions with disrupted directional coherence. Moreover, we illustrated the conceptual basis for these tensor-valued metrics (**Figure 5B**). MK_I_ reflected diffusion restrictions in all directions, typically arising from swollen or densely packed cells, while MK_A_ captured diffusion anisotropy associated with aligned microstructures such as neurites.

**Figure 5.**
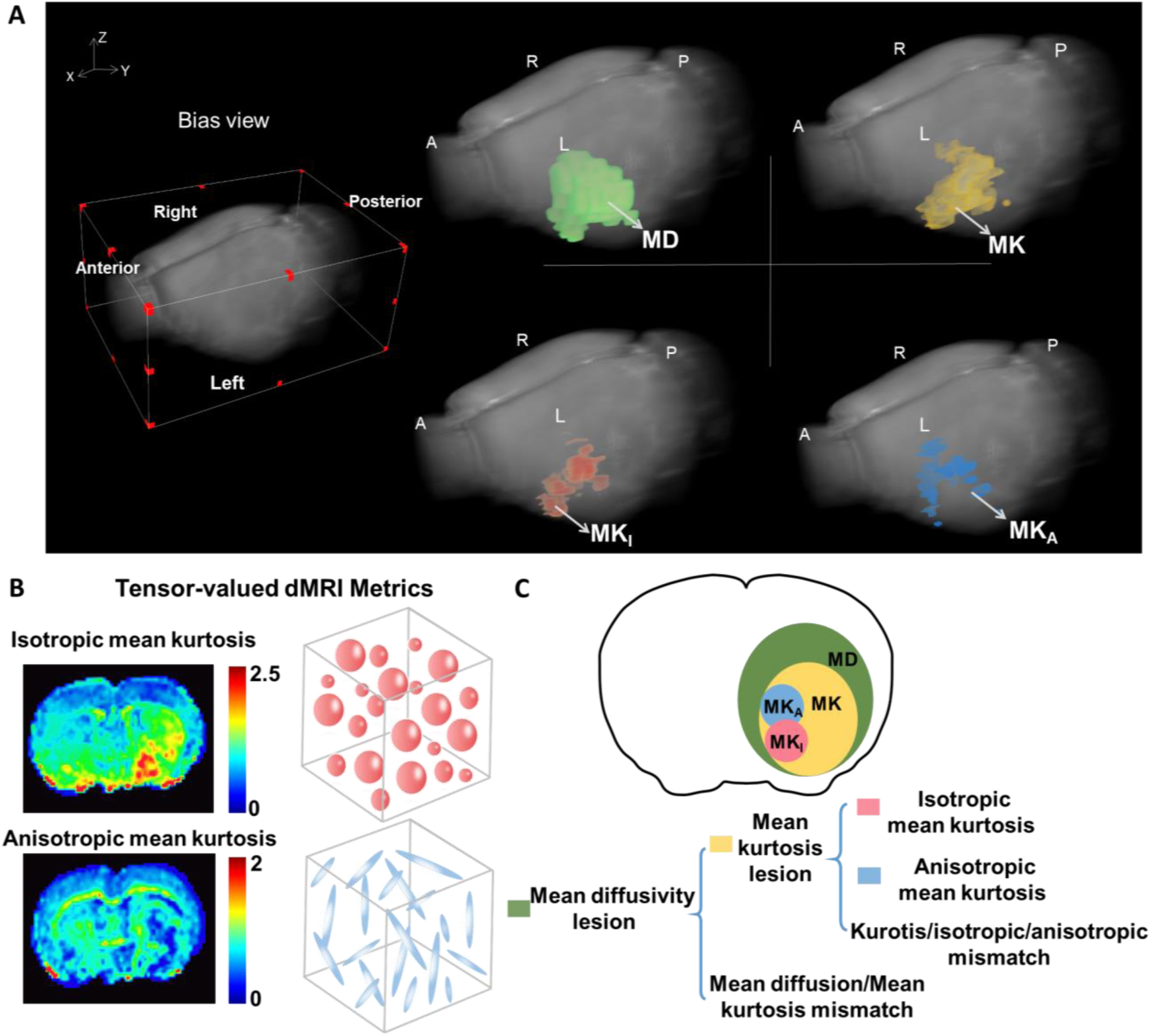
Tensor-valued diffusion MRI reveals mismatches in isotropic and anisotropic kurtosis components. **A**, Three-dimensional rendering of lesion volumes from a representative animal 2 hours post-MCAO, segmented by diffusion and kurtosis metrics: mean diffusivity (MD, green), mean kurtosis (MK, yellow), isotropic kurtosis (MK_I_, red), and anisotropic kurtosis (MK_A_, blue). **B**, Schematic representations of isotropic (spherical) and anisotropic (columnar) microstructural environments, corresponding to MK_I_ and MK_A_ components. **C**, A conceptual framework illustrating how Tensor-valued dMRI extends traditional lesion characterization. While MD identifies the infarct core, kurtosis imaging refines this into distinct regions based on MK_I_ and MK_A_ profiles, revealing mismatches between diffusion loss and microstructural kurtosis alterations.

Together, we inferred potential mismatch regions by examining the spatial overlap and divergence among MD, MK, MK_I_, and MK_A_ maps **(Figure 5C)**. Although the lesion masks overlapped, they did not fully coincide, suggesting that each metric captures distinct microstructural features. We hypothesize that regions with reduced MD but normal MK_I_ may reflect penumbral tissue with intact cellular density but impaired diffusion, while areas with high MK_I_ and preserved MK_A_ may indicate dense but structurally coherent tissue. By integrating these multidimensional contrasts, we proposed a novel framework to further delineate the infarct core, complementing existing paradigms such as perfusion-diffusion and diffusion-kurtosis mismatch.

## 4. Discussion

In this study, we applied tensor-valued diffusion MRI to a rat model of acute ischemic stroke, enabling in vivo decomposition of mean kurtosis (MK) into isotropic (MK_I_) and anisotropic (MK_A_) components. This methodological advance provided enhanced sensitivity and specificity in characterizing the heterogeneous microstructural alterations associated with acute cerebral ischemia. Our findings demonstrate that MK_I_ and MK_A_ not only differ in their spatial distribution within the infarcted brain, but also correlate strongly with histological indices of cellular density variations and fiber organization, respectively. These results suggest that tensor-valued dMRI may offer a more mechanistic and tissue-specific approach to delineating the infarct core and characterizing its internal microstructural heterogeneity, surpassing the capabilities of conventional DTI or standard DKI.

Consistent with previous studies using DWI and DKI, we observed significant reductions in mean diffusivity (MD) and elevations in mean kurtosis (MK) within the ischemic territory, reflecting restricted water mobility and increased tissue heterogeneity. By separating MK into MK_I_ and MK_A_, we were able to distinguish spherically restricted diffusion likely attributable to cytotoxic edema (captured by MK_I_) from disruption of oriented structures such as axonal fibers and dendrites (reflected by MK_A_). Elevated MK_I_ was primarily localized to the infarct core, correlating with histological evidence of cytotoxic edema and cell density variations^29^. In contrast, MK_A_ was significantly reduced in areas where myelinated fiber organization was disrupted, as validated by Luxol Fast Blue staining. Both our work and previous studies have shown that H_A_ is proportional to the MK_A_ derived from the macroscopic structural tensor obtained through histological analysis^30^. The ability to decouple these diffusion components allowed us to map both spherically restricted and directionally impaired compartments, enhancing specificity beyond that of DKI.

Our findings are in agreement with a recent human study^31^, which applied b-tensor diffusion encoding in acute stroke patients. Their work demonstrated that diffusional variance metrics, including MK_A_ and MK_I_, were sensitive to ischemic changes and modulated by diffusion time. Specifically, longer diffusion times revealed elevated microscopic fractional anisotropy (μFA) and MK_A_ in lesions compared to contralateral regions, whereas shorter diffusion times yielded reduced values. The detectability of these structural changes as either anisotropic or isotropic signals depends critically on the diffusion time window. In our animal study, diffusion data were acquired at a single diffusion time during the subacute phase, a period likely characterized by fully developed cytotoxic edema and substantial axonal damage. This may explain the predominance of isotropic kurtosis elevation and anisotropic kurtosis reduction observed in the infarct core. Despite differences in timing and protocol, both studies suggest that tensor-valued diffusion MRI can disentangle isotropic and anisotropic tissue features. This capability highlights the technique’s potential for clinical translation, offering a more specific and biologically meaningful assessment of stroke pathology than conventional diffusion methods.

Importantly, our findings suggest that an anisotropic-isotropic kurtosis mismatch may serve as a novel biomarker to refine the delineation of infarct core in ischemic stroke. This mismatch reflects underlying microstructural heterogeneity and implies that kurtosis decomposition can differentiate pathophysiologically distinct compartments within the lesion. Specifically, elevated MK_I_ coupled with reduced MK_A_ may correspond to regions of irreversible damage, whereas areas with preserved MK_A_ but mild MK_I_ elevation may represent functionally impaired yet structurally viable tissue. As visualized in our 3D lesion reconstructions, MD, MK, MK_I_, and MK_A_-defined lesions suggests the presence of multiple ischemic tissue subtypes. These may include infarct core, salvageable penumbra, and metabolically stressed but structurally preserved zones that remain indistinct on conventional diffusion imaging. From a translational perspective, tensor-valued dMRI offers a biologically meaningful framework for interpreting lesion complexity, and may complement or enhance existing paradigms, such as the perfusion-diffusion mismatch used in clinical stroke imaging, by adding microstructural specificity. This approach could ultimately support more personalized treatment decisions in acute stroke management.

Despite these promising findings, several limitations should be acknowledged. First, the relatively small sample size (n=6) may limit generalizability, although inter-animal consistency was high. Second, the study was performed at ultra-high field (9.4T) with small voxel size, which may not directly translate to clinical settings; however, tensor-valued diffusion encoding is feasible at clinical field strengths and has been demonstrated in human studies^31^. Third, while histology provides ground truth, co-registration between MRI and histological sections remains technically challenging and may introduce minor spatial misalignments^28, 32^.

Future research should aim to investigate the temporal dynamics of mismatch, tracking their evolution across different stages of stroke. Such longitudinal analysis could offer critical insights into the progression, severity, and potential reversibility of tissue injury^33^. In addition, extending this approach to other stroke models, including hemorrhagic stroke and various ischemic subtypes, would help elucidate both shared and distinct patterns of microstructural disruption. These efforts would enhance the clinical applicability and generalizability of tensor-valued diffusion MRI as a versatile tool for characterizing diverse forms of cerebrovascular injury.

## 5. Conclusion

We have demonstrated that in vivo tensor-valued diffusion MRI can separately quantify isotropic and anisotropic diffusion kurtosis in acute stroke, revealing a distinct kurtosis mismatch in ischemic tissue that is not discernible with conventional techniques. To our knowledge, this is the first application of tensor-valued dMRI in an experimental stroke setting, and it establishes anisotropic-isotropic kurtosis mismatch as a novel imaging marker of microstructural damage. Compared to standard DTI and DKI, the tensor-valued method offers enhanced specificity and has the potential to refine the identification of the injured core versus salvageable tissue. These findings contribute to the growing evidence that advanced diffusion MRI can capture the complexity of ischemic tissue heterogeneity.

In summary, tensor-valued diffusion MRI deciphers kurtosis to be isotropic and anisotropic kurtosis, thereby potentially enhancing sensitive microstructural readouts in stroke. The ability to map anisotropic and isotropic kurtosis in vivo opens new possibilities for stroke imaging: from improving early diagnosis and core delineation to monitoring therapies that target edema or cytoskeletal integrity. Ultimately, this work underscores the potential of advanced diffusion MRI not just to detect stroke, but to decode its underlying microstructural signatures, paving the way for better prognostication and targeted interventions in stroke care.

## Acknowledgments

This work was supported by grants from the National Natural Science Foundation of China (No. 82201447, No. RLZY20231001-01, and No. 82371307), the Strategic Priority Research Program of the Chinese Academy of Sciences (XDB0930000), Shenzhen Science and Technology Program (JSGGKQTD20210831174329010), National Key Research and Development Program of China (2023YFF0714200, 2024YFC3406602), the Chinese Academy of Sciences Project ( No. Y2021098), the CAS Project for Young Scientists in Basic Research (YSBR-114), the Fundamental Research Funds for the Central Universities (YG2025ZD18, YT), Young Leading Scientists Cultivation Plan supported by Shanghai Municipal Education Commission (ZXWH1082101, YT) and Key Laboratory for Magnetic Resonance and Multimodality Imaging of Guangdong Province (2023B1212060052), the Medical Scientific Research Foundation of Guangdong Province of China (A2023080).

## CRediT authorship contribution statement

**Mingyao Liang:** Methodology, Formal analysis, Investigation, Visualization, Writing - original draft, Writing - review & editing. **Jianyu Yuan**: Investigation, Methodology, Visualization, Writing - review & editing. **Chaogang Tang**: Investigation, Methodology, Visualization, Writing - review & editing. **Pingfu Wang**: Investigation, Methodology, Visualization, Writing - review & editing. **Tingting Gu**: Investigation, Methodology, Writing - review & editing**. Zhao-Yu Deng**: Investigation, Methodology, Writing - review & editing. **Weiming Sun**: Methodology, Investigation, Writing - review & editing. **Xinming Yang:** Investigation, Methodology, Writing - review & editing. **Yaohui Tang**: Methodology, Investigation, Writing - review & editing. **Ye Li:** Investigation, Methodology, Resources, Writing - review & editing. **Yi He**: Conceptualization, Resources, Investigation, Visualization, Writing - original draft, Writing - review & editing.

## Nonstandard Abbreviations and Acronyms

MRI: magnetic resonance imaging
MCAO: middle cerebral artery occlusion
DWI: diffusion weighted imaging
DKI: diffusion kurtosis imaging
MD: mean diffusivity
MK: mean kurtosis
MK_A_: anisotropic mean kurtosis
MK_I_: isotropic mean kurtosis
MK_T_: total mean kurtosis
H_A_: histological anisotropy
H_I_: histological isotropy

